# Temperature conditions allowing metamorphosis of *Hyalomma marginatum* nymphs: insights from laboratory experiments

**DOI:** 10.64898/2026.10.18.760010

**Authors:** Frédéric Stachurski, Maxime Duhayon, Anne-Sophie Guin, Carla Giupponi, Madiou Thierno Bah

## Abstract

Since the establishment of the *Hyalomma marginatum* tick, one of the main vectors of the Crimean-Congo haemorrhagic fever virus, has been recorded in mainland France, populations have only been observed in areas characterized by a Mediterranean climate. During the lifecycle of this two-host species, only two stages, the eggs laid by engorged females and the engorged nymphs, develop in the environment. Studies were carried out to understand the climatic factors, mainly the temperature (T°), which restrict its spreading. The moulting of the engorged nymphs was thus studied in climatic chambers at different constant T° (18°C to 40°C) and under various daily fluctuating T° cycles (the coldest was 8-20°C and the warmest was 16-30°C). The mortality rate of the engorged nymphs was lower than 4%, except for the lowest tested T° (18°C, 8-20°C and 8-22°C). Engorged nymphs could survive for a long time at 2°C (the mortality rate exceeded 10% only after 40 days at that T°) if the ticks were placed at that cold temperature immediately. On the other hand, they were more sensitive if they had already begun to develop before being placed at 2°C: when the metamorphosis had already started 8 days ago, the mortality rate reached 96% for the nymphs transferred for 3 weeks at cold T°. Using different methods, the development threshold was determined to be around 14°C. The median moulting time depended on the number of degree-days (D°) accumulated daily over the threshold. It ranged from 10 days (at 40°C) to 68 days (at 18°C) and was shorter for the T° cycles than for the constant T° equal to the mean T° of the cycles (68 days at 18°C but 52 days at 14-22°C and 37 days at 10-26°C). Consequently, the mean daily temperature is inadequate to assess the number of D° experienced by the moulting ticks. Formulas are proposed to determine the number of daily D°. The moulting required 196 D° (median time) for nymphs placed at fluctuating T° and 237 D° for those exposed to constant T°. It is suggested that this is due to inertia in nymph development: metamorphosis does not stop immediately when the T° goes back under the threshold.

## Introduction

The establishment of *Hyalomma marginatum*, one of the main vectors of the Crimean-Congo haemorrhagic fever (CCHF) virus in Eurasia (Bente et al., 2013), has been recorded in the French mainland for about a decade (Vial et al., 2016; Bah et al., 2022). This tick is a two-host species (Walker et al., 2003): engorged larvae moult while still attached to the host, and the resulting unfed nymphs immediately reattach to the same host. Therefore, only two stages of development occur off-host: egg-laying of engorged females followed by incubation and hatching of the eggs, and moulting of the engorged nymphs.

The phenology of *H. marginatum* was studied over several years in a southern France site where the species was abundant and where a horse herd grazed freely 24 hours a day on a 200-ha land (Stachurski et al., 2026). Adult ticks infested horses mainly between March and July-August, with an infestation peak in May. Immature ticks parasitized birds, their main hosts in this site, from mid-June to late September. Consequently, females likely lay their eggs from spring onwards, whereas nymph moulting takes place during summer. Experiments carried out under quasi-natural conditions in the same site revealed that nymphs moult within 3-4 weeks in July-August. In contrast, the nymphs that detached after mid-September mostly died before completing their metamorphosis.

There is clear evidence that engorged *H. marginatum* nymphs spread by migratory birds into several northern European countries can moult into active adults if environmental conditions are favourable. Such cases have been recorded, among others, in Germany, the Netherlands and Sweden (Chitimia-Dobler et al., 2019; Grandi et al., 2020; Uiterwijk et al., 2021; Chitimia-Dobler et al., 2024). However, to date, no populations seem to have established following these introduction events.

The development of *H. marginatum* nymphs is temperature-dependent, with the duration of moulting varying according to the temperature of exposition. As with insects (Gullan and Cranston, 2014), a temperature threshold allowing metamorphosis can be determined and the number of required “degree days” (D°) for moulting, accumulated above the threshold, can be estimated. Models based on “degree days” accumulation are widely used in entomology and plant pathology to predict pests and vectors activity (Murray, 2020). Determining this threshold for *H. marginatum* would enable the identification of regions where introduced nymphs could successfully moult, as well as an estimation of the time required for this process.

According to Gale et al. (2012), moulting of *H. marginatum* nymphs requires temperatures above 8°C for 15 consecutive days. Gray et al. (2009) quoted that Emelyanova reported that about 300 cumulative degrees above the developmental zero (14-16°C) are needed to complete the moult. These criteria lead to different conclusions, raising questions about the conditions responsible for the moulting of *H. marginatum* nymphs in populations established in southern France. Although the relationship between temperature and *H. marginatum* nymph moulting time is well established (Ouhelli, 1994; Elhachimi et al., 2021), the precise temperature threshold above which development occurs remains undetermined. In addition, engorged nymphs developing in the field are exposed to fluctuating temperatures, sometimes varying by as much as 15-20°C daily. Such fluctuations can affect insects’ development period in comparison with constant temperature conditions, either speeding up or slowing down the process, depending on the species and the temperature range (Hagstrum and Milliken, 1991; Paaijmans et al., 2010; Manenti et al., 2021). If a similar phenomenon occurs with ticks, the mean temperature of a region may not adequately reflect *H. marginatum* nymphs moulting kinetics.

To address these issues, metamorphosis of *H. marginatum* nymphs was studied between 2019 and 2024 under controlled conditions at different temperatures. The results were then compared to field observations previously reported (Stachurski et al., 2026).

## Material and methods

### Breeding of H. marginatum

In 2018, 17 *H. marginatum* engorged females, collected on horses from the herd monitored since 2016 (Stachurski et al., 2026), were placed for oviposition in the Baillarguet insectarium platform (https://doi.org/10.18167/infrastructure/00001). Part of the eggs and the females recovered before death were sent to the “Laboratoire de santé animale” of the “Agence nationale de sécurité sanitaire de l’alimentation, de l’environnement et du travail” (ANSES), Maisons-Alfort, to look for the CCHF virus using two different RT-qPCR assays (Wölfel et al., 2007; Sas et al., 2018). All ticks and egg batches were CCHF-free. The remaining eggs were therefore kept to initiate a breeding colony of the species. This breeding colony and the experiments described below were carried out in the insectarium platform. The breeding was set up according to the procedures described below, authorized by the French Ministry of Research (No. APAFIS#6304-2016080315594914 v4 of 05 Feb. 2018 and APAFIS#43847-202306161342860 v5 of 24 Oct. 2023 for « Feeding of ticks on vertebrates for laboratory breeding »).

Unfed adult ticks were placed in cloth bags glued to the ears of goats kept in an animal house. Each bag, closed with rubber bands, contained 5-7 males and an equal number of females. The bags were opened daily to check for tick attachment and to observe female feeding. If ticks died during the first two days, new specimens of the same sex were placed in the bags. After detachment, engorged females were placed in a climatic chamber (Memmert HPP110) to lay eggs. The eggs were separated into batches of 500-800 eggs (about 25-40 mg) and regularly monitored until hatching.

One- to three-month-old larvae were placed in cloth bags glued on the back of New Zealand rabbits (Charles River, L’Arbresle, France), maintained at 20-21°C. Nymph detachment was checked starting from day 12 after larvae installation. Engorged nymphs were collected daily and placed in climatic chambers.

### Development of engorged nymphs in climatic chambers

Groups of at least 100 nymphs were subjected to 12 different constant temperature conditions (T_c_), ranging from 18°C to 40°C, with a relative humidity (RH) of 85% (Table 1). Sinusoidal daily fluctuating temperatures (T_f_) were also programmed into the climatic chambers. Ten different programmes were tested, some sharing the same mean value (Table 2). The objective was to replicate the temperature variations observed at different times of the year in the site where *H. marginatum* infestation had been monitored. The coldest cycle (8-20°C, mean: 14°C) matched the temperatures observed in the latter half of April. The warmest cycle (16-30°C, mean: 23°C) was slightly cooler than the temperatures recorded in July-August, when engorged nymphs detach from their bird hosts (Stachurski et al., 2026). The range between minimum and maximum temperatures varied from 8 to 16°C, depending on the cycle. Nymph metamorphosis was monitored once daily, at approximately the same time each day.

**Table 1.** Moulting time of *Hyalomma marginatum* engorged nymphs at different constant temperatures.

| Temperature °C |  | 18°C | 19°C | 20°C | 21°C | 22°C | 23.5°C | 25°C | 27°C | 28°C | 32°C | 36°C | 40°C |
| --- | --- | --- | --- | --- | --- | --- | --- | --- | --- | --- | --- | --- | --- |
| Number of nymphs |  | 472 | 323 | 576 | 180 | 349 | 242 | 605 | 101 | 1333 | 115 | 218 | 182 |
| Moulting time (days) | Median | 68 | 44 | 41 | 36 | 29 | 25 | 20 | 18 | 17 | 13 | 11 | 10 |
|  | Mean | 82.5 | 45.5 | 43.0 | 35.9 | 28.5 | 25.2 | 19.7 | 18.4 | 17.0 | 13.0 | 10.7 | 9.8 |

**Table 2.** Moulting time of *Hyalomma marginatum* engorged nymphs at different fluctuating temperatures.

| Extremes of the tested temperature cycles |  | 8°-20°C | 8°-22°C | 10°-22°C | 10°-24°C | 14°-22°C | 12°-24°C | 10°-26°C | 15°-26°C | 13°-28°C | 16°-30°C |
| --- | --- | --- | --- | --- | --- | --- | --- | --- | --- | --- | --- |
| Mean temperature of the cycle |  | 14°C | 15°C | 16°C | 17°C | 18°C | 18°C | 18°C | 20.5°C | 20.5°C | 23°C |
| Number of nymphs |  | 113 | 268 | 354 | 210 | 248 | 205 | 168 | 204 | 128 | 193 |
| Moulting time (days) | Median | 96 | 72 | 57 | 44 | 52 | 42 | 37 | 31 | 31 | 25 |
|  | Mean | 98.2 | 74.5 | 58.7 | 44.1 | 52.7 | 42.7 | 36.9 | 31.6 | 31.5 | 24.6 |

To assess the effect of thermal regime on nymph development, one constant-temperature regime (18°C) was compared with three fluctuating regimes sharing the same mean temperature but increasing amplitudes (14-22°C, 12-24°C, and 10-26°C). The development rate (i.e., the reciprocal of the moulting time) was analysed using a linear model with thermal regime as the explanatory variable.

The nymphs were kept in complete darkness, except for one trial carried out to determine whether light affected moulting time. Three different climatic chambers were used for this trial, implemented at 28°C. In one chamber, nymphs were always kept in constant darkness (24N). In the second, they were in darkness for 16 hours (16N), with light on (2000 lux) for the remaining 8 hours. In the last chamber, the light was on for 16 hours (2000 lux) and darkness lasted for 8 hours (8N). Two trials were conducted, with each of the three lighting conditions successively programmed in two different climatic chambers. The effect of lighting conditions on moulting time was analysed using a linear mixed effects model, with light regime (8N, 16N, 24N) as a fixed effect and random intercepts specified for each trial and for the climatic chamber nested within trial. For both analyses (effect of lighting regime and of climatic chamber), estimated marginal means and pairwise comparisons were computed using the marginal effects package (Arel-Bundock et al., 2024) with P-values adjusted for multiple comparisons using Tukey’s method.

Finally, batches of engorged nymphs were placed at 2°C for different periods and then transferred to 25°C in order to determine whether the ticks were still alive and if metamorphosis could occur. The RH in the climatic chamber (ESPEC SH-241) could not exceed 40% at this low temperature. Fourteen batches of 30-80 ticks were placed at 2°C immediately after nymph detachment for periods ranging from 8 to 120 days. Eight other batches of 19-28 ticks were placed at 2°C after 4, 8, 12, and 16 days at 25°C, periods during which the nymphs had started their moulting process. They were transferred back to 25°C after one or three weeks. The goal was to compare the mortality between the nymphs placed at low temperature straight after their drop-off and those which had already begun their metamorphosis before being moved to 2°C.

### Determination of the temperature threshold (T_t_)

Various methods exist to assess this value. The simplest approach is the one described for insects by Gullan and Cranston (2014). After establishing the average moulting time required at different constant temperatures, the reciprocal of these times (i.e., the development rate) is plotted against the temperature. A linear regression line is then fitted to these points, allowing for the calculation of the line’s slope and the threshold temperature, which is the X value when Y=0 (the developmental zero).

The variance in development time varied with temperature, indicating heteroskedasticity. To account for it, quantile regression was used which, unlike the least square method, models the conditional means of the response variable and estimates the effects of the predictor variable across different points of that conditional distribution. This approach allowed explicit expression of the changes in the shape of the distribution of the response variable (development time) across the predictor variable (temperature), highlighting the temperature threshold for each conditional distribution of the tick populations. The chosen quantiles were the median (q50), q10, q25, q75 and q90, the highest development rates (q90) being observed among the nymphs with the shortest moulting times. Models fitting was done using the *nlme* package (v3.1.166; Pinheiro and Bates, 2000) and the quantreg package (v5.98; Koenker, 2005) for the generalized least square model, the mixed-effect model and the quantile regression respectively. All analyses were conducted with R version 4.3.2 (R Core Team, 2023).

### Determination of the number of degree-days (D°)

The number of D° above the threshold T_t_ that permits moulting can be determined using the linear regression line equation. It can also be easily calculated for the trials carried out at any given constant temperature (T_c_), with each day contributing T_c_ – T_t_ degree-days. To determine the number of D° accumulated with fluctuating temperatures (T_f_), it was necessary to retrieve the actual temperature evolution in the climatic chambers, recorded minute by minute. This made it possible to determine the number of degree-minutes, then degree-hours, and finally D° for each tested programme.

### Assessment of the number of D° in the field

To determine the number of D° daily accumulated by moulting nymphs in the field, it is not always possible to have temperature records on a minute-by-minute or even an hourly basis. Typically, only the mean (T_m_), maximum (T_x_), and minimum (T_n_) daily temperatures are available. The simplest approach is to calculate the difference between the daily mean temperature and the threshold temperature using equation A:

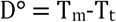

Considering that mean temperature is inadequate, Gardiner et al. (1981) proposed formulas to calculate the development rate for each half-day using maximal and minimal daily temperatures, assuming the temperature changes follow a quasi-sinusoidal curve. Similar to Gardiner, sine-wave methods (Baskerville and Emin, 1969) assumed that daily temperature variation adheres to a sine-shaped curve and that daily effective temperature is equal to the area under the curve given a temperature threshold T_t_. The corresponding equation (B) is the following:

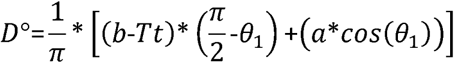

with a = (T_x_ - T_n_)/2, b = (T_x_ + T_n_)/2 and θ_1_ = sin^-1^[(T_t_ - b)/a]

Wine growers use the Huglin index to classify a region’s climate and determine suitable vine types (Bonnefoy et al., 2010). It is calculated as follows:

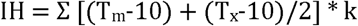

Both mean and maximum temperatures are compared with 10°C, considered as the development threshold temperature for the vine, and k is a coefficient linked to the latitude. According to this model, different proxies, using only T_m_, T_n_ and T_x_ were compared to the D° determined for the different T_f_ as explained above. The best fit was obtained with the equation C:

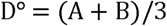

with A = (T_m_-T_t_) if T_m_ > T_t_ or A = 0 if T_m_ < T_t_, and B = (T_x_-T_t_) if T_x_ > T_t_ or B = 0 if T_x_ < T_t_

### Validation of the calculated values by field observations

To verify whether the calculations made using data collected under controlled conditions were consistent with the field observations, the study carried out in 2019-2020 was revisited. Engorged nymphs had been placed at different times, between July and October (4 cohorts), in cages installed in the *garrigue* (Stachurski et al., 2026). Cages were examined several weeks and months later to check for nymph metamorphosis. The temperature in the experimental enclosure was recorded on an hourly basis throughout the study. This allowed the determination of the number of D° accumulated for each study period. It was then possible to verify whether the metamorphosis kinetics was compatible with the calculations made using the laboratory data.

## Results

### Breeding of H. marginatum

Between 2018 and 2025, 10,581 *H. marginatum* nymphs were collected. On average, they weighed 24 ± 0.002 mg (12-53 mg) and detached from their hosts after 19.6 ± 0.0005 days (13-34 days) of feeding. More than 7,600 of these nymphs were involved in the studies presented below.

### Mortality rate

The mortality rate of the nymphs placed at constant temperatures was lower than 4%, except for those installed at 18°C which reached 18.6% (Figure 1 and Data D1). The mortality rate of the nymphs exposed to fluctuating temperatures was also less than 4%, except for the two cycles with the lowest means and a minimum temperature of 8°C (8-22°C and 8-20°C), with mortality rates of 14% and 27%, respectively (Figure 1 and Data D1). Some of the dead ticks in the three trials with a high mortality rate (18°C, 8-20°C, 8-22°C) were adults unable to emerge from their thicker than usual cuticle. When removed from the exuviae, these adults were dead and poorly formed (twisted legs, articles shorter than normal, deformed joints).

**Figure 1.**
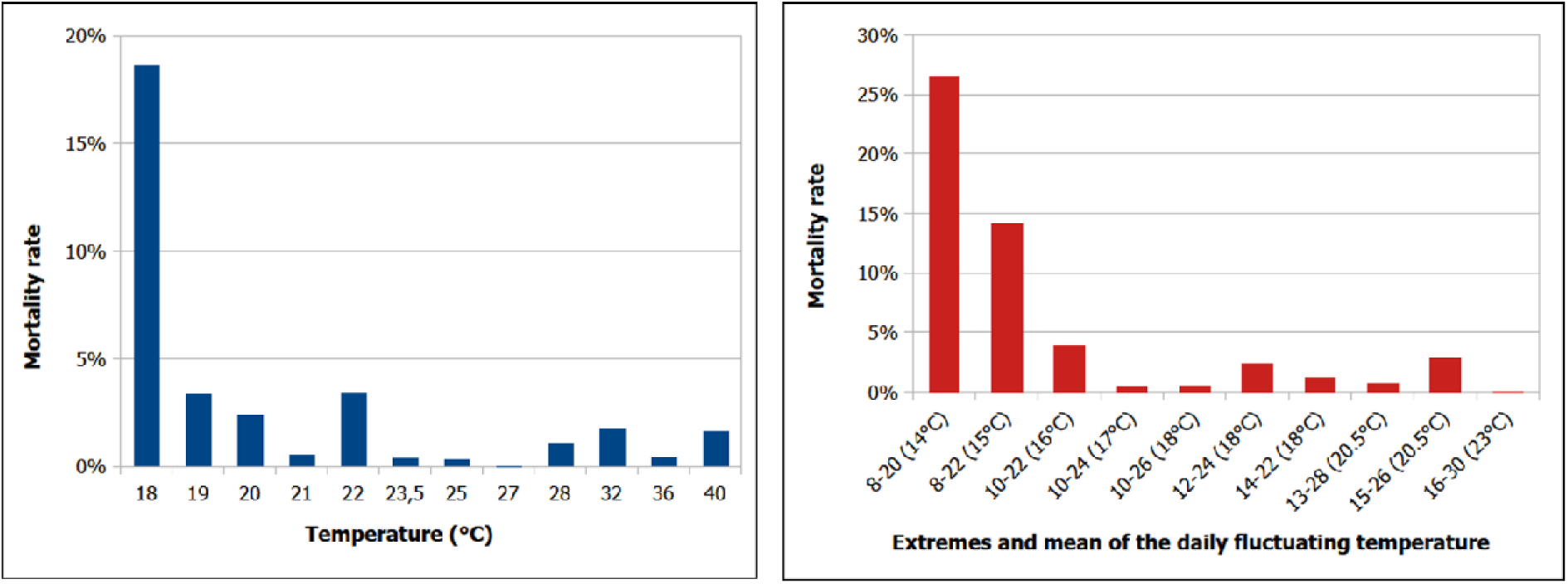
Mortality rate of engorged *Hyalomma marginatum* nymphs placed at different constant (left graph) or fluctuating (right graph) temperatures.

### Effect of cold temperature on the mortality rate

The mean mortality rate of the nymphs placed at 2°C immediately after detachment was 6% (0-13%) for the nine batches left in the cold for a maximum of 40 days. It increased to about 50% after 60 or 75 days at 2°C and finally to 90% for the nymphs exposed three or four months at this cold temperature (Figure 2 and Data D2).

**Figure 2.**
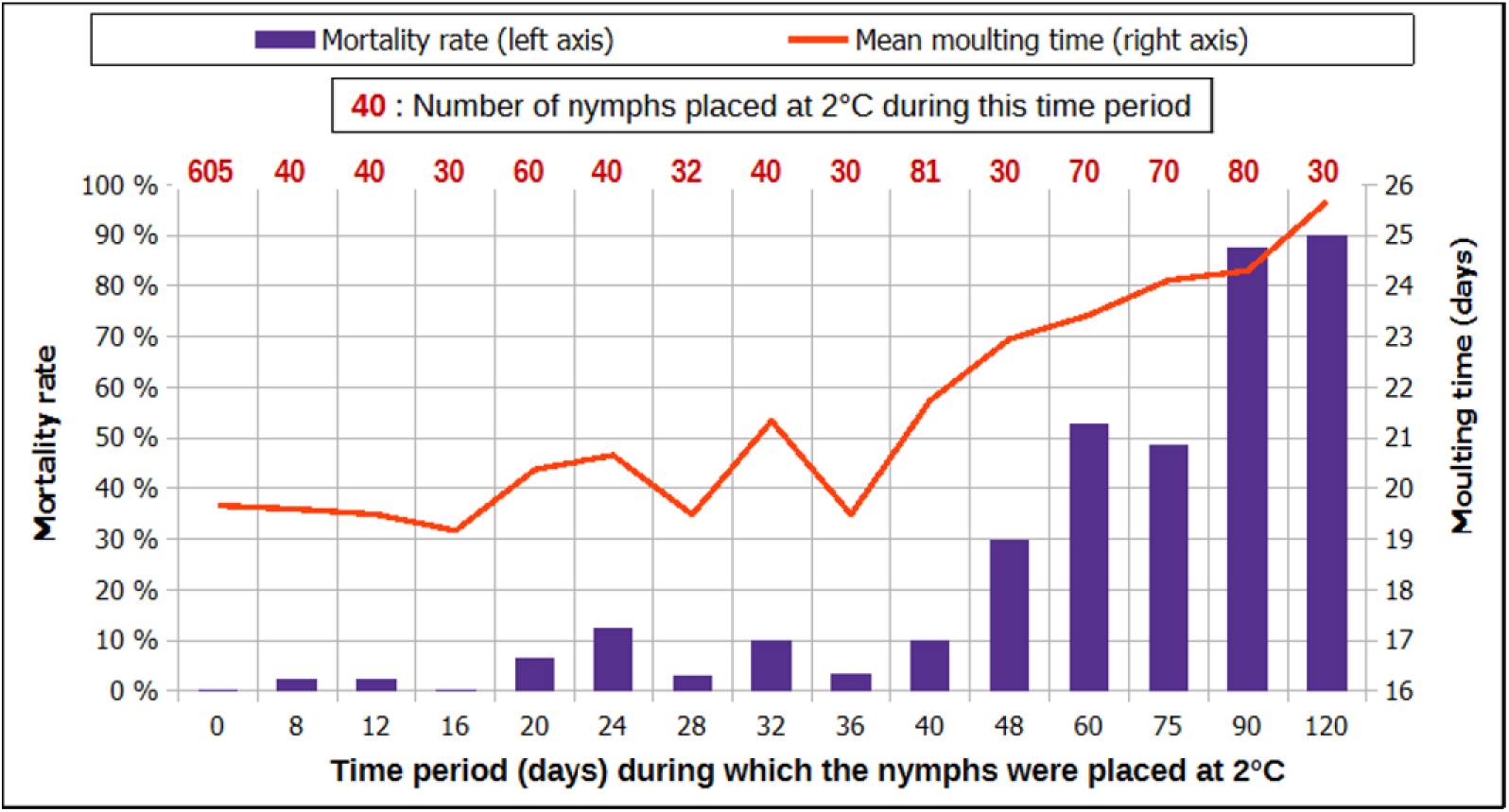
Mortality rate and mean moulting time of engorged *Hyalomma marginatum* nymphs placed at 2°C for different time periods immediately after detachment.

Nymphs placed at 2°C while they had started their development at 25°C displayed higher mortality rates (Figure 3 and Data D2).

**Figure 3.**
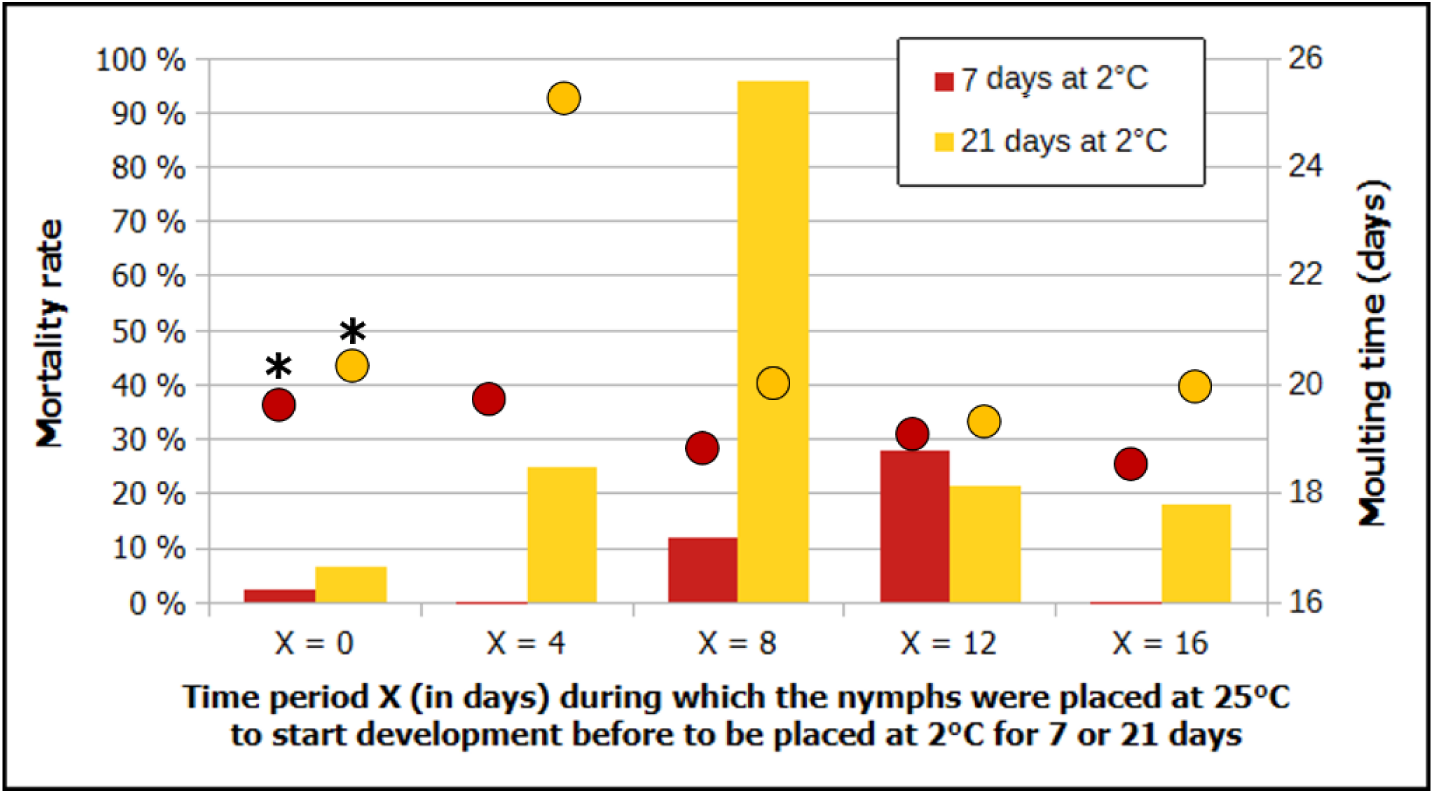
Mortality rate (bars, left axis) and mean moulting time (coloured dots, right axis) of engorged *Hyalomma marginatum* nymphs placed at 2°C for one or three weeks while they had started their development at 25°C for various time periods. Results observed during preceding trials (see figures 1 and 2) are marked by a star.

The mean mortality rate of the batches placed at cold for 3 weeks after the development had started was 40% (18-96%) whereas it was 7% and 13% for the nymphs exposed to 2°C for 20 or 24 days, respectively, immediately after drop-off (Figure 2). The mortality rate was 12% and 28%, respectively, for the nymphs placed at 2°C for one week after 8 or 12 days of development at 25°C, but only 3% if they were transferred immediately to 2°C for 8 days. No death was recorded in the batches placed in the cold for one week, 4 and 16 days after the beginning of metamorphosis. The highest mortality rate (96%) was noted for the nymphs transferred to 2°C for 3 weeks, whereas metamorphosis had started 8 days prior.

### Moulting time of *H. marginatum* nymphs

The development time gradually decreased with increasing temperature for the ticks placed at T_c_. The median moulting time was 68 days at 18°C or 41 days at 20°C, but 20 days at 25°C and 10 days at 40°C (Table 1). This corresponds to a development rate of 0.015, 0.024, 0.050 and 0.100, respectively. The variability of development duration was higher for the low temperatures (Figure 4 and Data D1).

**Figure 4.**
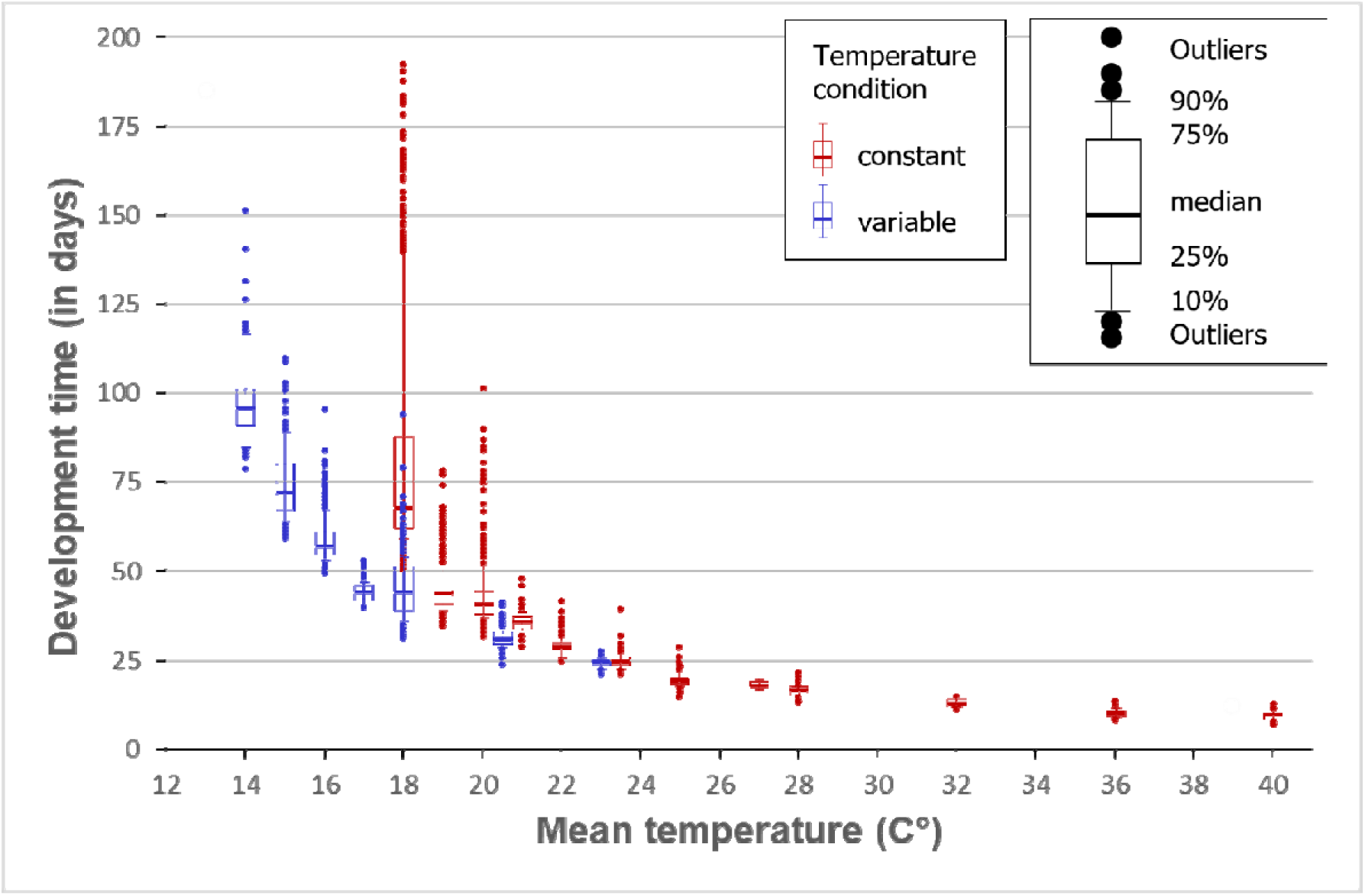
Development time of *Hyalomma marginatum* nymphs placed at different constant and fluctuating temperatures. Note that the data for the three cycles with a mean of 18°C and the two cycles with a mean of 20.5°C are grouped together.

The same pattern was observed with T_f_. The median moulting time was 96 days for the cycle 8-22°C (mean: 14°C) and 25 days for the cycle 16-30°C (mean: 23°C) (Table 2 and Figure 4). The moulting time was shorter for the nymphs placed at T_f_ than for those placed at the constant temperature corresponding to the mean of the fluctuating cycle. The median moulting time was thus 31 days for the cycles 13-28°C and 15-26°C (mean temperature: 20.5°C), whereas it was 41 days at 20°C and 36 days at 21°C.

It appeared that a wider thermal amplitude was associated with accelerated development for an identical mean temperature. Pairwise comparisons showed that all regimes differed significantly from one another (p < 0.001, Table 3). While the median moulting time at 18°C was 68 days, it was 52 days for the 14-22°C cycle, 42 days for 12-24°C and 37 days for 10-26°C.

**Table 3.** Estimated marginal means, standard error and confidence interval the of development rate under four temperature regimes sharing the same mean temperature (18°C). Means not sharing a letter differ significantly (Tukey-adjusted pairwise comparisons, α = 0.05).

| Temperature regime | Amplitude (C°) | Estimated mean of the development rate | Standard Error | 95% CI | Group |
| --- | --- | --- | --- | --- | --- |
| 18°C (constant) | 0°C | 0.0134 | 0.00029 | 0.0129 – 0.0140 | a |
| 14-22 °C | 8°C | 0.0191 | 0.00036 | 0.0184 – 0.0198 | b |
| 12-24°C | 12°C | 0.0235 | 0.00040 | 0.0228 – 0.0243 | c |
| 10-26°C | 16°C | 0.0272 | 0.00044 | 0.0263 – 0.0281 | d |

### Effect of cold temperature on the moulting time

The mean moulting time of the nymphs placed back at 25°C after several days or weeks at 2°C fluctuated between 19.2 and 21.7 days, with a slight upward trend for the batches maintained in the cold up to 40 days after their detachment (figure 2). It then increased up to 25.7 for the few nymphs that successfully moulted despite 1.5 to 4 months at 2°C. The mean moulting time of the nymphs placed at 2°C 4-16 days after beginning metamorphosis was more or less similar (18.5-20 days) to that of the nymphs placed at 25°C immediately after drop off (19.7 days), except for the batch put at 2°C for three weeks, four days after detachment. In that case, the moulting time reached 25.3 days (Figure 3).

### Effect of light on the moulting time

The mean moulting time for the nymphs kept at 28°C either in complete darkness (24N) or with light for 16 or 8 hours of the nychthemeron (16N and 8N), was 16.9 days. This was shorter than the time observed during a previous study, carried out a year before with 74 nymphs kept in complete darkness (17.6 days). More precisely, the mean moulting time was 16.2, 16.8 and 16.7 during the first trial, under the conditions 24N, 16N and 8N, respectively, whereas it was 17.3, 16.7 and 17,0 days for the same three lighting conditions during the second trial. The linear mixed-effects model detected no significant effect of the pair lighting regime-climatic chamber on moulting time (F = 0.03, p = 0.971). None of the pairwise contrast approached significance after Tukey adjustment (Supplementary Material S1).

### Determination of the developmental temperature threshold (T_t_)

When assessed by the linear regression of the mean development rate at different constant temperatures plotted against the temperature, the T_t_ was 13.90°C. When estimated from quantile regression, it ranged from 13.75 to 14.67 (Table 5, Figure 5). The different methods therefore converged on a similar developmental threshold, which was set to 14 °C for subsequent calculations.

**Table 5.** Parameter estimates and 95% confidence interval (CI) of the linear functions y = ax + b for the different quantile regression models. The respective temperature thresholds were calculated by the formula -b/a and the required degree-days (D°) for the metamorphosis by the formula 1/a.

| Quantile | Slope (coefficient a) | | | Intercept (coefficient b) | | | Assessed threshold | Number of $D^\circ$ |
| --- | --- | --- | --- | --- | --- | --- | --- | --- |
|  | Estimate | 95%CI | p-value | Estimate | 95%CI | p-value |  |  |
| q10 | 0.00395 | 0.00385 – 0.00433 | <0.001 | -0.0579 | -0.0632 – -0.0551 | <0.001 | 14.67 | 253 |
| q25 | 0.00402 | 0.00377 – 0.00453 | <0.001 | -0.0569 | -0.0704 – -0.0509 | <0.001 | 14.17 | 249 |
| q50 | 0.00422 | 0.00402 – 0.00435 | <0.001 | -0.0592 | -0.0620 – -0.0529 | <0.001 | 14.05 | 237 |
| q75 | 0.00442 | 0.00409 – 0.00474 | <0.001 | -0.0612 | -0.0712 – -0.0519 | <0.001 | 13.86 | 226 |
| q90 | 0.00468 | 0.00430 – 0.00524 | <0.001 | -0.0643 | -0.0720 – -0.0459 | <0.001 | 13.75 | 214 |

**Figure 5.**
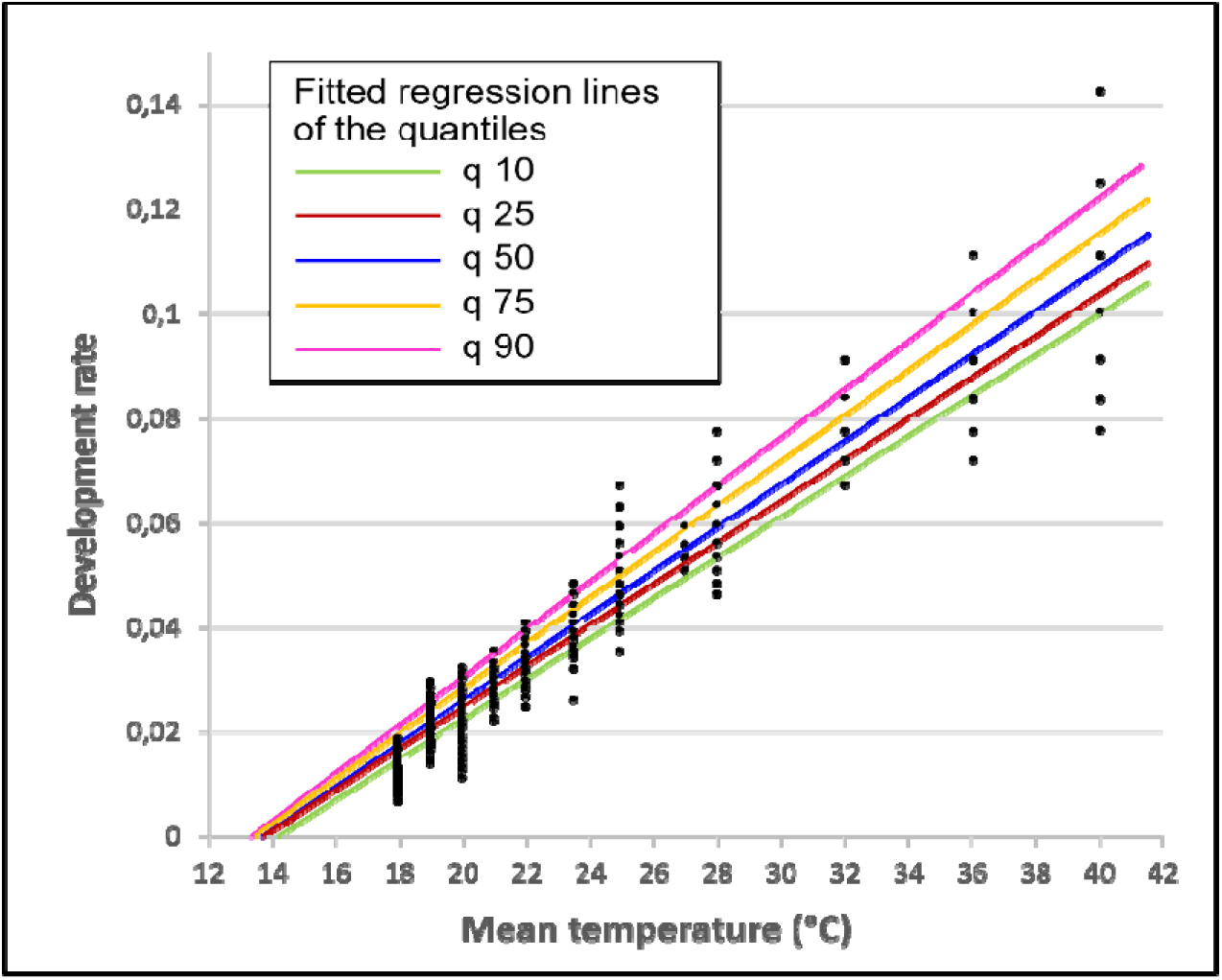
Quantile regression lines of development rate (i.e., the reciprocal of the moulting time) of *Hyalomma marginatum* engorged nymphs according to the temperature. Fitted lines show the 10^th^, 25^th^ ,50^th^ ,75^th^ and 90^th^ conditional quantile (q10-q90).

### Determination of the number of degree-days (D°) required for nymph moulting

The number of D° above the development threshold required for nymph moulting can be determined by multiplying the development time (in days, see Table 1) by the value T_c_-T_t_, the threshold T_t_ being 14°C. When the mean moulting times are considered, metamorphosis occurred after accumulating 330 D° at 18°C, 217 D° at 25°C or 255 D° at 40°C. The average number of D° for the 12 constant temperatures tested is 246 D°. This value can also be calculated using the regression line equations, as the reciprocal of the slopes (Table 5). It varies from 214 D° for q90 (the nymphs with the highest development rate, i.e., the shortest moulting time) to 253 D° for q10, being 237 D° for the median. Lastly, the number of D° required for metamorphosis can be assessed with the results from the trials carried out with T_f_, as the D° accumulated daily by the nymphs in development could be determined precisely (see Material and Methods, Table 6 and Data D3). The median moulting time ranged from 176 to 226 D° (mean: 196 D°). As for the above assessment, it increased with the proportion of nymphs considered: 181 D° for the first decile, 188 D° for the first quartile, 220 D° for the ninth decile (Table 6).

**Table 6.**
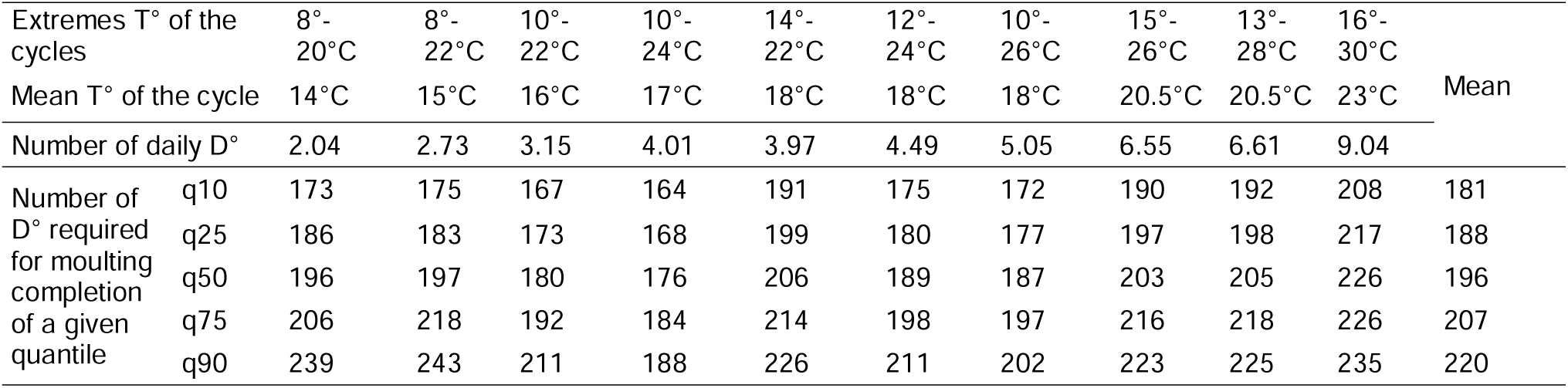
Number of degree-days (D°) allowing the metamorphosis of various proportion of engorged *Hyalomma marginatum* nymphs placed at different fluctuating temperature (T°).

### Assessment of the number of D° in field conditions

There was a good correlation (r = 0.984, t = 15.65) between the actual number of D° accumulated daily with the various fluctuating cycles and the D° calculated with equation A, which only took into account the mean daily temperature (Supplementary Material S2). The correlation was better (r = 0.991, t = 21.45) with equation C in which D° is estimated with the mean and maximal daily temperature (S2). As equation A fits better with the real D° for temperature cycles with a high mean, and equation C fits better with temperature cycles with a low mean, a combination of these two equations was used to assess the daily D° (S2). Equation A was used when minimal daily temperature was higher than the threshold temperature (14°C), while equation C was used in the opposite case. With that combination, the correlation was higher (r = 0.998, t = 43.14). When equation B was used in combination with equation A when the minimal daily temperature was higher than the threshold temperature, the correlation reached nearly 100% (r = 0.999, t = 5654.4). However, for computational simplicity, the combination of equations A and C was used for the calculations presented in the following section.

### Validation of the calculated values with field observations

During the field study carried out in 2019-2020, it was observed that all ticks from the first cohort recovered three weeks after the nymphs that had been installed in the cages had moulted (Figure 6). A total of 235 D°, calculated with the combination of equations A and C (Data D4), had accumulated at that time. In the first cage examined from cohort 2, 21 of the 24 recovered ticks (88%) were recently-moulted adults. A total of 195 D° had accumulated when this cage was examined. The three unmoulted nymphs were placed at 25°C in the insectarium, of which one died and the two others moulted in less than one day (i.e., when less than 11 additional D° were provided). Three weeks later, when 321 D° accumulated in total, all the recovered nymphs were inactive adults.

**Figure 6.**
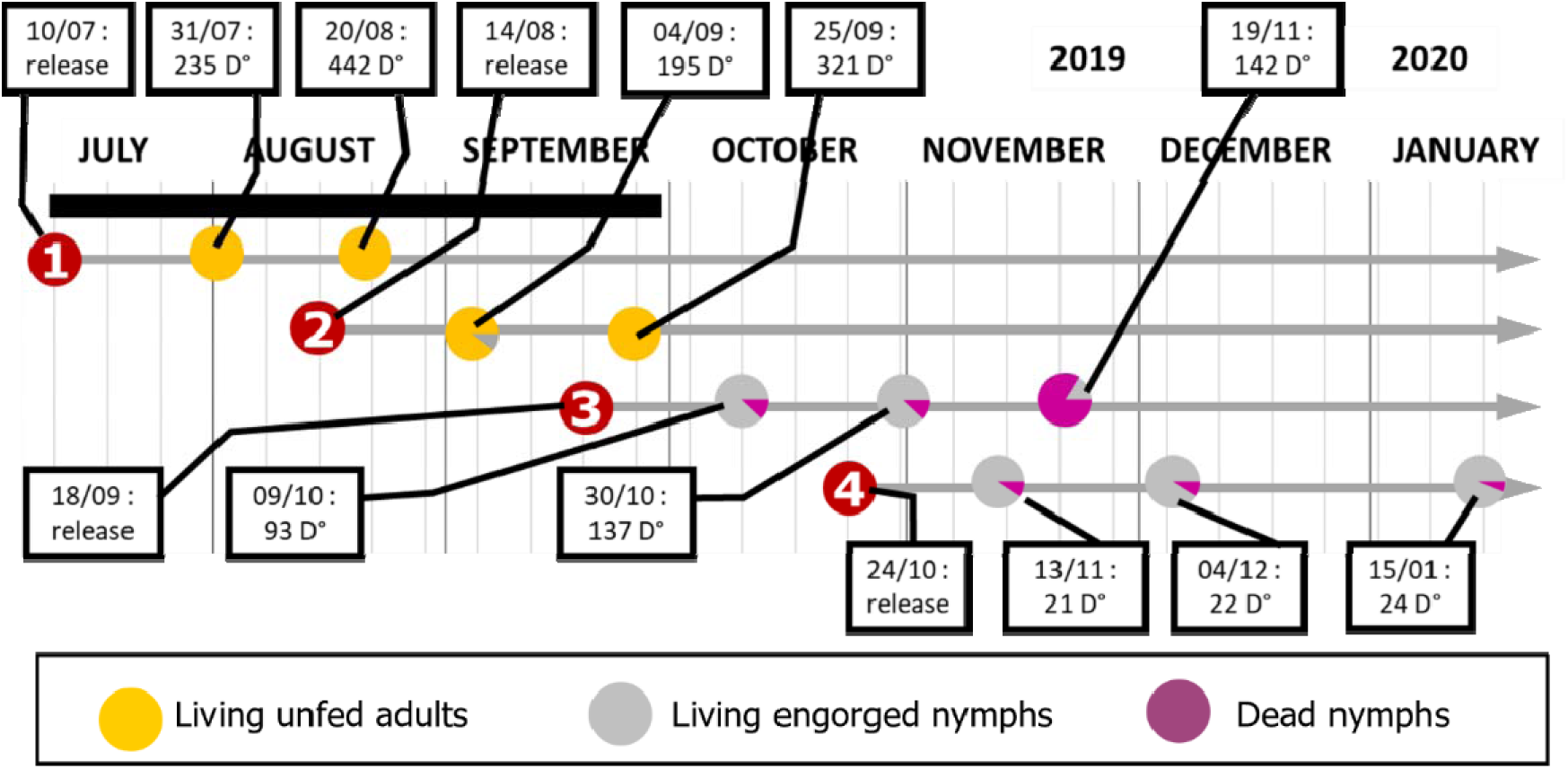
Graphic representation of the status of the ticks found in the cages where batches of 26-30 engorged nymphs had been placed. Four cohorts were studied. The predated nymphs are not taken into account. The unmoulted nymphs were placed at 25°C and 85% RH in a climatic chamber to verify whether they were still alive. The number of degree-days (D°) accumulated while the different cages were examined was calculated using the meteorological data recorded on the study site and the equation A (D° = T_m_-T_t_) when the minimal temperature was higher than the threshold temperature (T_t_) or the equation C (D° = [(T_m_-T_t_) + (T_x_-T_t_)]/3) in the opposite case (T_m_: mean temperature; T_x_: maximal temperature). The black line indicates the period of natural nymph detachment from their bird hosts (for more details, see Stachurski and al., 2026).

The situation was completely different with cohorts 3 and 4: no adults were recovered from the cages examined between October and January. A total of 93, 137 and 142 D° accumulated, respectively, when the three cages of cohort 3 were examined (Figure 6). When put back at 25°C in the insectarium, the live nymphs required 11, 6 and 5 days (median value), respectively, to moult (i.e., 121, 66 and 55 D°). In total, the median moulting time of the nymphs from cohort 3 was about 214, 203 and 197 D° for the three successive cages. The nymphs from cohort 4 had accumulated only a few degree-days (21, 22 and 24 D°, respectively) when they were recovered. Most of them were still mobile and required 16 to 26 days at 25°C to moult (median time: 18 days, i.e., 198 D°). In total, the median moulting time of the nymphs from this cohort was about 220 D°.

Meteorological data collected during the study period showed that the climate was not more severe during the 3 weeks preceding the examination of cage 3 from cohort 3 (44 mm rainfall; mean temperature: 8.7°C; minimal temperature -0.7°C; mean minimal temperature during the period: 5.8°C), than during the 3 weeks preceding the examination of cages 2 and 3 from cohort 4 (respectively, 162 mm rainfall; 7.6°C; -0.7°C; 4.6°C and 67 mm rainfall; 8.1°C; -1.5°C; 4.2°C). The mortality rate of the nymphs in cage 3 from cohort 3 was, however, greatly higher. It increased from 15% or 18% (cages 1 and 2 examined on 09/10/2019 and 30/10/2019, respectively) to 83% for cage 3 examined on 19/11/2019. On the contrary, the mortality rate of the non-predated nymphs from cohort 4 remained low during the same period: it was 10% in cage 1 (examined on 13/11/2019), 8% in cage 2 (04/12/2019) and 6% in cage 3 (15/01/2020).

## Discussion

Within the range of the tested temperatures, metamorphosis of *H. marginatum* nymphs occurred logically more quickly at higher temperatures. The moulting times observed during the present study were generally close to those recorded during other studies carried out in Morocco. At 25°C, Ouhelli (1994) observed a mean moulting time of 18.6 to 20.4 days, depending of the RH, whereas it was 19.7 days in the current study. Elhachimi et al. (2021) reported a median moulting time of 68 days at 18°C, as in the present study, and 19 days at 28°C, compared to 17 days in this study. The greatest difference was noted at high temperature: Ouhelli (1994) recorded a mean moulting time of 4 days at 35°C, while, in this study, it was 10.7 days (median time: 11 days) at 36°C.

Depending on the method used and the proportion of nymphs considered, the temperature threshold required for metamorphosis in *H. marginatum* nymphs was assessed to be between 13.8°C and 14.7°C. In addition to the natural idiosyncrasy of the ticks, even when maintained at the same temperature, some biases have affected the determination of the moulting times and, consequently, these calculations. First, nymphs were collected only once per day, and may have started their development before being placed in the climatic chambers, although the rabbits were reared at 20-21°C. Second, nymphs were fed on 17 different rabbits, which were more or less tolerant of tick infestation, and their weight after feeding ranged from 12 to 53 mg. Both factors may have impacted the development time. Third, as metamorphosis was recorded only once per day, moulting could have occurred anywhere from a few minutes to almost 24 hours before being observed. Lastly, five different climatic chambers were used, and their settings were not completely identical, as shown by the results obtained with the three lighting conditions tested in three different climatic chambers: the moulting times depended as much on the chamber as on the regime. Each of these inaccuracies corresponds to a maximum of approximately 10 D°, except for the once-daily record of metamorphosis, which may represent an overestimation of the moulting time of up to 22 or 26 D° for the highest temperatures (36°C and 40°C). A perfect determination of the threshold is therefore impossible. It has consequently been set at 14 °C, which roughly corresponds to the average of the various calculation methods. It is not known whether this threshold is a consistent characteristic of all *H. marginatum* populations, or whether it might vary genetically between geographically distinct populations.

The moulting times at fluctuating temperatures were always shorter than those observed when the nymphs were placed at a constant temperature equivalent to the mean of the tested cycles. Moreover, moulting times were not similar for the three cycles with a mean temperature of 18°C. This shows that the daily mean temperature does not allow for a correct assessment of the number of degree-days accumulated each day. This is confirmed by the fact that moulting could occur for nymphs placed at 8-20°C, a cycle for which the mean was similar to the temperature development threshold. The apparent little importance of the daily mean temperature, provided that daily maximum temperatures exceed the threshold, could be particularly advantageous for the establishment of *H. marginatum* in continental-type environments, characterised by strong daily thermal amplitudes, where a low mean temperature would otherwise seem unfavourable for development.

According to the different quantiles, the number of D° needed for nymph metamorphosis with fluctuating temperatures was 33-42 D° lower than that needed with constant temperatures. This suggests some inertia in nymph development: metamorphosis does not slow down or stop immediately when the temperature decreases below the threshold. This would explain why the number of D° accumulated at metamorphosis was lower for the 10-26°C cycle than for the 12-24°C or 14-22°C cycles. With that latter cycle, the temperature was never below the threshold, so the possible inertia effect could not fully operate. The results of the first three cages from cohort 3 also reinforce this hypothesis: the nymphs that were exposed to the natural fluctuations of the temperature in the field for longer periods before being placed at constant temperature in the insectarium needed, overall, fewer D° to moult. The use of formula C, combining mean and maximal daily temperatures instead of only mean temperature, to assess the daily number of accumulated D° when the daily minimal temperature is lower than the threshold, makes it possible to take the inertia of development into account. Field observations validated the laboratory results overall.

*Hyalomma marginatum* nymph moulting occurred with high success across a large temperature spectrum. No trial was carried out to determine the maximum constant temperature tolerated by *H. marginatum* engorged nymphs. The mortality rate ranged from 0 to 4% for the tick batches exposed to a constant temperature between 19°C and 40°C or to fluctuating temperatures with the lower limit of at least 10°C. It increased drastically when they were placed at 18°C, the lowest constant temperature tested, or when the lower bound of the cycle was 8°C. The nymphs also survived a long time when placed at 2°C immediately after detachment from their rabbit host. The mortality rate exceeded 50% only after two months at this low temperature. The nymphs were far more sensitive to cold when they had already started their development: the mortality rate reached 96% for the nymphs kept at 2°C for 3 weeks, after having developed for 8 days at 25°C and having accumulated 88 D° before experiencing the cold. This increased sensitivity of developing nymphs may explain the high mortality of nymphs placed at 8-20°C and 8-22°C. It could also explain the difference observed between cohorts 3 and 4 during the field trial. The mortality rate was 83% for nymphs from cohort 3, whereas it remained low (6-10%) for ticks from cohort 4, which were subjected to the same meteorological conditions. Ticks from cohort 3 had already accumulated about 140 D° when the cold occurred, whereas those from cohort 4 had not yet clearly started metamorphosis at that moment, having only accumulated 21-24 D°. This differential cold tolerance, depending on the degree of progression during the moulting process, highlights the importance for nymphs to complete metamorphosis within a sufficiently short time window once development has started. The length of this window probably depends on local environmental conditions: in Morocco or Spain, for instance, nymphs may still moult readily in September, when temperatures remained warm, whereas the same period is already largely unsuitable in southern France.

Some of the nymphs that died when placed at 18°C, 8-20°C or 8-22°C showed the same deformities and inability to emerge from their exuviae as some of the ticks from cohorts 3 and 4 during the field trial (Stachurski et al., 2026). A long stay in adverse conditions seems to prevent ticks from completing metamorphosis successfully. However, in the field trials, it was not possible to determine whether the cold was the sole cause, or whether rain could have also caused damages to nymphs that were already undergoing metamorphosis, and were therefore unable to move and escape the adverse conditions.

The number of D° accumulated when the metamorphosis occurred varied between nymphs, attesting to their idiosyncratic behaviour. This amount varied from 153 D° to 370 D° for the 1992 nymphs which moulted at fluctuating temperature (only 1% of the nymphs needed more than 300 D° to moult). Ouhelli (1994) recorded a mean moulting time of 115.5 days at 16°C, corresponding to 231 D°, which is consistent with the results obtained in the present study at constant temperatures. Conversely, he recorded a mean moulting time of 4 days at 35°C, i.e., 84 D°, which is significantly lower than all the moulting times observed during the present study, even with fluctuating temperatures. The first decile of the surveyed nymphs needed 153-170 D° to moult and half moulted with less than 200 D°. This individual variability raises the question of whether a small number of ticks, able to moult with fewer degree-days or to tolerate colder conditions, could be sufficient to allow a population to establish in environments at the margin of *H. marginatum* favourability. This question is directly linked to the concept of effective population size. At the field study site in July and August, 50 D° accumulated in about 5 days, indicating that most of the nymphs disseminated at that period could moult. Conversely, after mid-September, the daily number of D° drastically decreased, as did the possibility of metamorphosis.

Two hundred D° over the 14°C threshold can easily be obtained anywhere in France, and probably across Europe, during the 4-5 warmest months of the year. For example, according to data collected in the regional airports by Météo-France, 370 D° accumulated in Lille (Northern France) between 01/07 and 31/08/2025, 419 D° in Strasbourg (Eastern France), 440 D° in Nantes (Western France) and 475 D° in Clermont-Ferrand (Central France) in comparison with 604 D° in the southern study site. Indeed, in a transplant experiment carried out in Gardouch (Southwestern France), engorged nymphs released in an open meadow in March–April successfully moulted into adults within 2 to 3 months, although these newly emerged adults remained mostly inactive (Giupponi et al., 2026). This may explain why engorged nymphs, disseminated by migrating birds, can moult in Germany, the Netherlands, and Sweden. Consequently, nymph metamorphosis does not appear to be the limiting factor for the establishment of *H. marginatum* populations, provided that the larvae are present in the field sufficiently early, before summer.

One question, however, remains to be answered: what proportion/quantity of nymphs must successfully complete their metamorphosis to enable a population of *H. marginatum* to become established in a given area? Alternatively, it may be argued that metamorphosis is unlikely to be strongly limiting as long as larvae attach early enough in the season. This conclusion naturally leads to the question of egg-laying and incubation times, which will be examined in a companion article.

## Author contributions

FS conceived and organized the various laboratory studies. MD programmed the climatic chambers. FS, MD, ASG and CG produced the engorged nymphs and recorded tick moulting and death. FS designed the experimental “cages” system and carried it out with MD. FS prepared the data which were analysed by FS, MTB and CG. FS wrote the first version of the manuscript, which was completed by MTB and revised by MD, CG and ASG. All authors approved the final version of the manuscript.

## Acknowledgements

The authors would like to thank the Baillarguet insectarium platform, where all the laboratory studies were performed, and the owner of the land where the field studies were carried out. Iyonna Zortman, a native English speaker, aided in correcting the language.

## Data and Supplementary Material availability

All data and supplementary materials will be available at https://doi.org/10.18167/DVN1/QMPS5H when the manuscript has been accepted for publication. The data of the dataverse are:

D1. H_marginatum_nymph_development_at_different_temperatures.csv. Fate (moulting time or death) of engorged nymphs placed at different constant or fluctuating temperatures.

D2. H_marginatum_nymph_development_when_placed_at_2°C.csv. Fate (moulting time or death) of engorged nymphs placed at 2°C immediately after detachment or after they started their development at 25°C.

D3. Temperature_evolution_during_fluctuating_cycles.csv. Minute par minute evolution of the temperature during the 10 fluctuating cycles tested in the climatic chambers.

D4. Temperature_records_on_field_study_site.csv. Temperatures recorded, between 01/07/2019 and 15/01/2020, on the study site where survival and development of Hyalomma marginatum nymphs was observed under natural conditions.

The supplementary materials are:

S1. Pairwise contrasts between lighting regimes for moulting time, estimated from a linear mixed-effects model with random intercepts for trial and chamber nested within trial.

S2. Linear regressions between the actual number of degree-days (D°) daily accumulated during the trials carried out at different fluctuating temperatures and the number of D° assessed by different equations.

