## Supplementary material S1 for "Temperature conditions allowing metamorphosis of *Hyalomma marginatum* nymphs: insights from laboratory experiments"

**Supplementary material 1.** Pairwise contrasts between lighting regimes for moulting time, estimated from a linear mixed-effects model with random intercepts for trial and chamber nested within trial (Tukey-adjusted pairwise comparisons, $\alpha=0.05$).

| Comparison | Estimate | Standard Error | 95% CI | t-value | p-value |
| --- | --- | --- | --- | --- | --- |
| 8N – 16N | 0.108 | 0.473 | -2.68 – 2.90 | 0.228 | 0.972 |
| 8N – 24N | 0.091 | 0.473 | -2.70 – 2.88 | 0.192 | 0.980 |
| 16N – 24N | 0.017 | 0.473 | -2.80 – 2.77 | -0.036 | 0.999 |
