## Supplementary material S2 for "Temperature conditions allowing metamorphosis of *Hyalomma marginatum* nymphs: insights from laboratory experiments"

**Supplementary material 2.** Correlation between the number of degree-days ($D^\circ$) daily accumulated during the trials carried out at different fluctuating temperatures, determined thanks to the records of the minute by minute temperature, and the number of $D^\circ$ assessed by different equations.

$T_t$: threshold temperature; $T_m$: mean temperature of the cycle; $T_x$: maximum temperature; $T_n$: minimum temperature;

$a = (T_x - T_n)/2$; $b = (T_x + T_n)/2$; $\theta_1 = \sin^{-1}[(T_t - b)/a]$

For equation C, $A = (T_m - T_t)$ if $T_m > T_t$ or $A = 0$ if $T_m < T_t$, and $B = (T_x - T_t)$ if $T_x > T_t$ or $B = 0$ if $T_x < T_t$

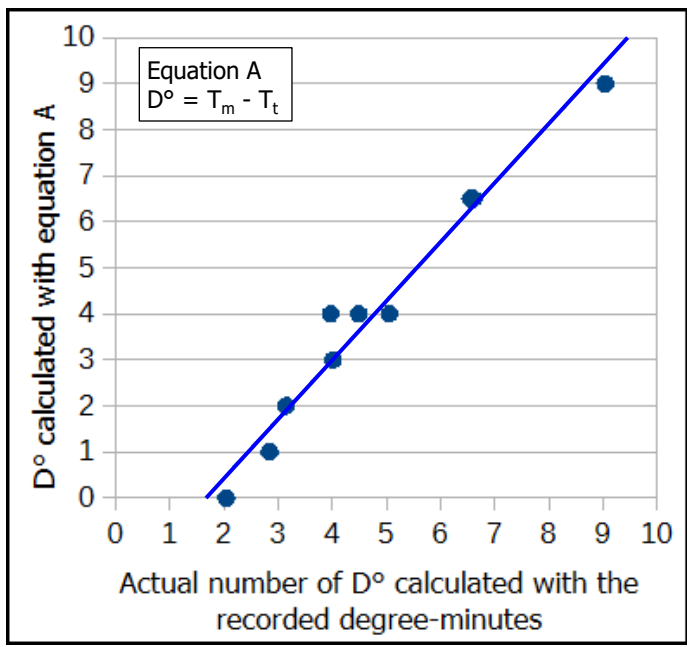

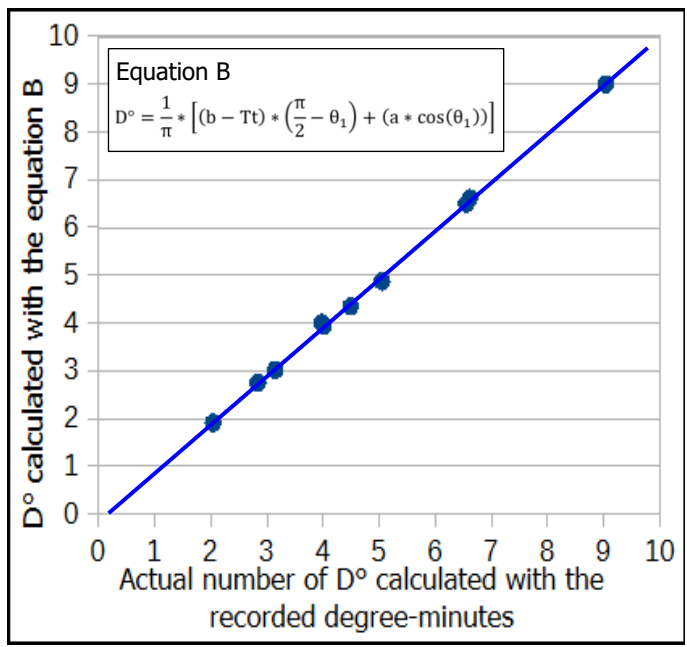

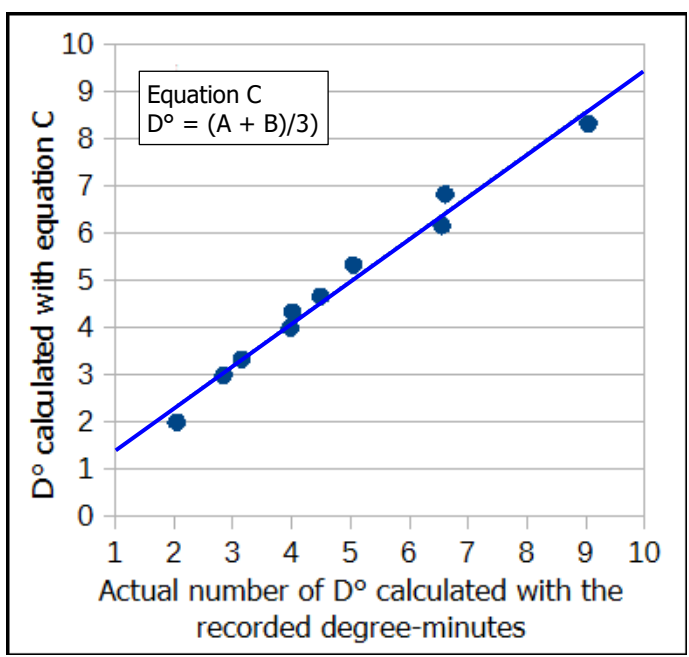

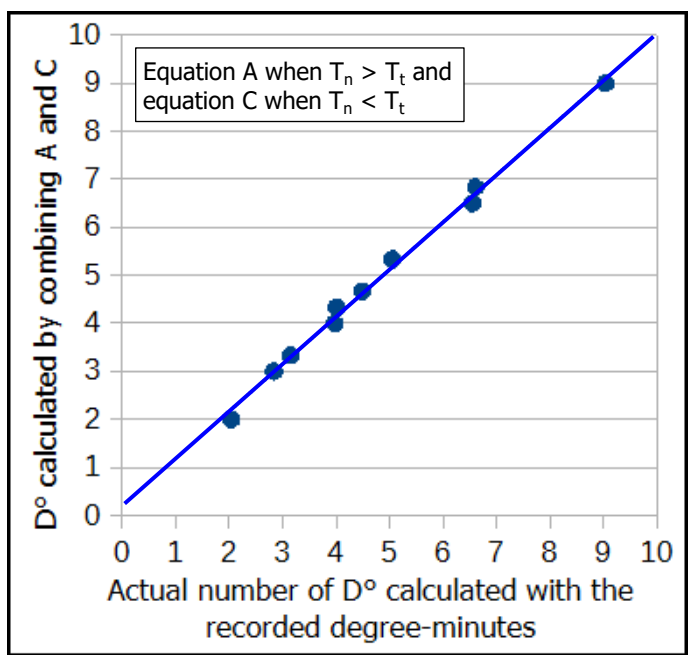
